# *Escherichia coli* antimicrobial resistance; Phenotypic and genotypic characterization

**DOI:** 10.1101/2021.12.06.471453

**Authors:** Nagwa Thabet Elsharawy, Hind A. A. Al-Zahrani, Amr A. El-Waseif

## Abstract

Improper use of the antimicrobials as *E. coli* giving the microorganism multi-resistance against many antimicrobials by gene mutation on integrons, transposons and plasmids. Therefore, our aim in this study is to 1)examine antibiotics resistance phenotype and genotype in *Escherichia coli*, 2) identifying the structure of bacterial resistance genes on whole-genome sequencing against multi-drug resistant of *Escherichia coli* in marketed poultry meat. Samples collected, prepared and Bacteriological examination, Antimicrobial sensitivity test performed, Serological identification of *Escherichia coli* isolates. Results declared that; the prevalence of *E. coli* from tested chicken meat samples of 100 chicken meat samples surveyed against *E. coli* the result declared that about 40%. Antimicrobial susceptibility was; antibiotics of choice against *E. coli* Sulfonamides, Cephalosporins,Tetracyclines, Quinolones. Serologically, STEC (O157:H7) 30%, ETEC (O142) 10%, EHEC (O26:H11). The subunit B of shiga-like toxin (SLT) gene appeared as a homogenous band. Heat-labile toxin (LT) gene was screened in both genomic DNA and plasmid preps in tested strains. Control STEC as it represents a danger to the poultry consumers. We recommended to increase the hygienic measures during slaughtering, processing and/or handling of chicken carcasses and avoidance unnecessary usage of any antimicrobials to avoid appearance of new antimicrobials resistant.

## Introduction

Although *E. coli* is a member of gram negative normal nonpathogenic intestinal inhabitant and consider as a commensal for animal and human being. It classified into four categories; enteroinvasive, enteropathogenic, enterohaemorrhagic or enterotoxigenic. However, drinking of contaminated water with or eating contaminated food can cause gastrointestinal diseases by pathogenic *E. coli*, (Shiga toxin-producing *E. coli)* (STEC), which appear as; haemorrhagic colitis (HC), diarrhoea and haemolytic-uraemic syndrome (HUS) [1].

STEC considered the more common pathogenic microbes transported via poultry meat to the human being and its commonly transmitted after consumption of dirty water, contaminated food such as; chicken, beef, sheep meat which mainly acquired contamination from fecal intestinal contents from the poultry itself during slaughter or from human handlers during storage and processing [2].

Antimicrobials used generally to treat any *E. coli* infection; the problem appear as the frequent improper use of the antimicrobials as *E. coli* therapy which giving the microorganism the multi-resistance against many antimicrobial drugs. The microorganism mainly has the antimicrobial resistance by gene mutation which leading to the presence of resistance genes on integrons, transposons and plasmids. Integron is the system used for gene expression and incorporate the cassettes of gene that activate it. Integron class I is comprising from; flanking a variable region (VR), 2 conversed segments (CS) the common form between Enterobacteriaceae new generations [3].

The antimicrobial resistance resulting from frequent antibiotics uses which leading to generating of new strains of microorganisms which become highly have resistance against several antibiotics. This considers one of the highly important crisis to the public health globally. Scientists looking for new generations of antibiotics to overcome the microorganism’s resistance in animals, poultry and human being [4].

Therefore, our aim in this study is to 1) examine antibiotics resistance phenotype and genotype in *Escherichia coli*, 2) identifying the structure of bacterial resistance genes on whole-genome sequencing against multi-drug resistant of *Escherichia coli* in marketed poultry meat.

## Material and methods

1. **Ethical approval:** There is no ethical approval necessary.
2. **Samples collection, preparation and Bacteriological examination:** the number of tested samples were about 100 chicken meat samples were collected randomly from different markets and stored in a polyethylene bag then immediately transferred inside ice box to the bacteriological laboratory for analysis. Two grams from homogenated chicken sample inoculated in MacConkey broth for about 18 hr./37°C. Then, streaked onto MacConkey agar (Oxoid) plates for about 24 hr./37°C. then the Rose pink colonies streaked onto Eosin Methylene Blue (Oxoid) and incubated for about 24hr./37°C. typical Escherichia coli morphology is a large colony, green metallic sheen with blue-black. *E. coli* colonies identified morphologically, microscopically and by biochemical tests kits (bioMerieux API, France) [5]. Further identification by Serotyping; using antisera sets (Denka Seiken Co., Japan) according to WHO, [6].
3. **Antimicrobial sensitivity test:** performed using disc diffusion technique by Muller-Hinton agar against 15 antibiotic discs as following; Gentamicin (30 µg/disk), Streptomycin (30 µg/disk), Ampicillin (10 µg/disk), Penicillin (10 µg/disk), Cefepime (30 µg/disk), Cefotaxime (30 µg/disk), Ciprofloxacin (30 µg/disk), Flumequine (30 µg/disk), Trimethoprim (30 µg/disk), Sulfamethoxazole (30 µg/disk), Tetracycline (30 µg/disk), Doxycycline (30 µg/disk) the results done according to Alderman & Smith, [7].
4. **Serological identification of *Escherichia coli* isolates [8]** by slide agglutination test using standard monovalent and polyvalent *E. coli* antisera sets for definition of the Enteropathogenic types as following;emulsified microbial colony by 2 drops of saline on a glass slide. Addition of a loopful antiserum, Agglutination performed, a further portion of the colony cultured on nutrient agar slant then incubated at 37°C/24 hours for testing mono-valent sera. Prepared suspension of the microbe in saline and performed the slide agglutination tests to identify the O-antigen.
5. **Extraction of Nucleic Acids:** DNA extracted by GeneJet genomic DNA purification kit (ThermoFisher Scientific, USA). Summarized as following; centrifuge picked bacterial colonies for 10 min./5000 ×g, resuspended the cell pellet in 180 μL of Digestion Solution (provided in the kit) then added 20 μL of Proteinase K then thoroughly mixed, incubated at 56°C in water bath for about 30 min with continuous shaking until complete lysis. Vortexing 20 μL of RNase solution after adding to the mixture, then incubated for 10 min./37°C. addition of 200 μL of Lysis solution to the mixture then vortexing 400 μL of 50% ethanol after adding it to the mixture. transferred lysated cells then purified and centrifuged for 1 min./6000 × g. washed column by 500 μL washing buffer (I and II) then centrifuged for 2 min at maximum speed until complete removal of ethanol.Stored the purified DNA at −20°C until use. **The plasmid preparation**; by GeneJet plasmid DNA miniprep kit (Thermofisher Scientific, USA). According to the manual; about 1-3 ml. of the grown culture put in 1.5 ml microcentrifuge tubes then centrifuged at 12,000xg/ 2 min. Re-suspend the pellet in 250μl in ice cold resuspension buffer supplied with the kit and mixed by inverting of the tube about 5-6 times. Tubes incubate for 5 min./37 °C. transferred the supernatant and centrifuged at 10,000xg/30 sec. and wash by 500μl buffer (supplied with the kit)and recentrifuged at 10,000xg/30 sec. elution of DNA plasmids using pre-warmed ddH2O 50μl then incubated for 3 min./37°C then recentrifuged by maximum speed (14,000xg)/30 sec. **Gene Amplification:** PCR reactions gene amplified by 1μl purified genetic substances (genomic DNA/ plasmid preps), 2.5μl MgCl2, 5μl buffer, 1μl of primer (listed in **table 1** below), 0.25μl of Taq Polymerase enzyme mix 0.5μl of dNTPs and complete volume to 25μl by free-nuclease water. Resolve PCR products using 0.5µg/ml ethidium bromide and agarose gel (1%), determination of the size of the resolved products by 100bp DNA ladder. Then run Gel at 80V/50 min. documentation bythe gel system (Biometra, Goettingen, Germany). **Extraction of DNA Fragments from Agarose Gel:** elution of DNA fragment using agarose gel by DNA extraction kit (Thermofisher Scientific, USA). Fragmentation performed under UV light then preserved into 1.5ml tube. Then, centrifuged at 13,000xg/2 min. Washed the columns by 700µl washing buffer, then centrifuged for 1 min./ 37oC. add 50µl of buffer to elute on the spin column filter, then kept for 1 min./37oC followed by centrifuged at 13,000xg/2 min.

**Table 1.**
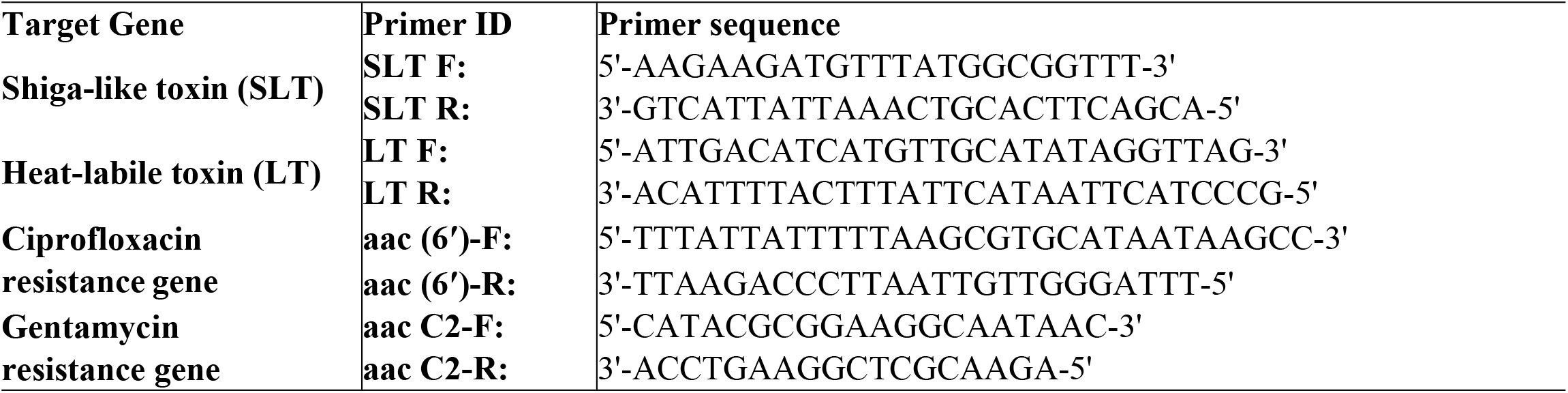
List of Primers Used in Gene Amplification

## Results

1. **Prevalence of *E. coli* from tested chicken meat samples:** According to (**fig 2)** A total of 100 chicken meat samples surveyed against *E. coli* the result declared that about 40% of the samples were positive while the rest samples had not *E. coli*.
2. **Antimicrobial resistance pattern for *E. coli* isolates:** The antimicrobial susceptibility testing of some *E. coli*strains (n=20) which isolated from chicken meat samples as illustrated in **table (2)** as following; antibiotics of choice against *E. coli* among 12 antimicrobial drugs related to five different classes were as following; the most effective antimicrobials related to **Sulfonamides**; trimethoprim (20/20) 100%, sulfamethoxazole (16/20) 80%, followed by **Cephalosporin’s** including; Cefepime (14/20) 70%, Cefotaxime (13/20) 65%, then **tetracycline’s** including; Tetracycline (10/20) 50%, Doxycycline (8/20) 40%, while among tested antimicrobial classes 3 family members recorded weak effect against *E. coli* as following; **Quinolones** including; Ciprofloxacin (4/20) 20%, Flumequine (4/20) 20%, **Aminoglycosides;** Gentamicin 3/20 (15%), Streptomycin (2/20) 10%, and the weakest members related to **β-lactame;** Penicillin 2/20 (10%), Ampicillin (zero%).
3. **Results for serological examination of *E. coli* isolates:** twenty *E. coli* isolates were serologically typed (**fig 3)**. Where 10/20 (50%) isolates were typed as STEC (O157:H7), 6/20 (30%) isolates were ETEC (O142), 2/20 (10%) isolate was EHEC (O26:H11). Finally, 2/20 (10%) of isolates were typed as EPEC (O55:H7). **Shiga-Like Toxin (SLT) Screening documented in (fig 4) as following:** the subunit B of shiga-like toxin (SLT) gene appeared as a homogenous band in their genomic DNA at 300 bp molecular weight. Results declared that strain (1) were the lowest amplification detected comparing to strain 2. **Heat-Labile Toxin (LT) Screening declared in (fig 5) as following:** Heat-labile toxin (LT) gene was screened in both genomic DNA and plasmid preps intested strains. A fragment of ∼ 200 bp was detected in both strains (1) and (2). **Gentamycin Resistance Screening observed in (fig 6) as following:** Gentamycin resistance gene (aac C2) fragment was detected in strains as afragment of molecular weight ∼ 856 bp but with a minor band of molecular weight of nearly 300 bp. **Ciprofloxacin Resistance Screening viewed in (fig 7) as following:** Ciprofloxacin-resistance gene was also screened in both genomic and plasmid preparations in strains under test. A sharp band at 1 kb was clearly detectedin strain (1) while it was absent in strain (2). In plasmid preps, no amplification of the target gene could be detectedin strains tested.
4. **Statistical Analyses:** analysis of variance will be conducted and means will compare using SPSS.

**Figure (1):**
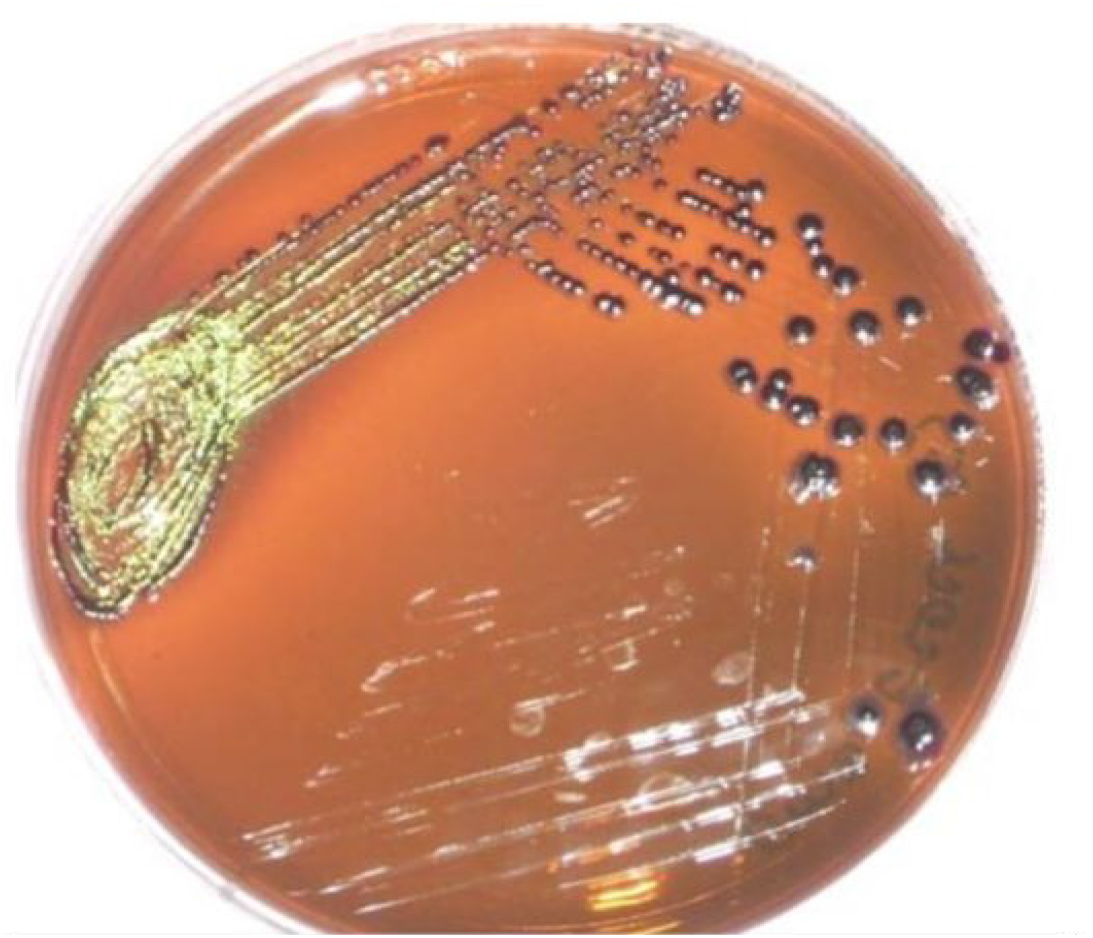
*E. coli* on eosin methylene blue (EMB) agar.

**Fig 2.**
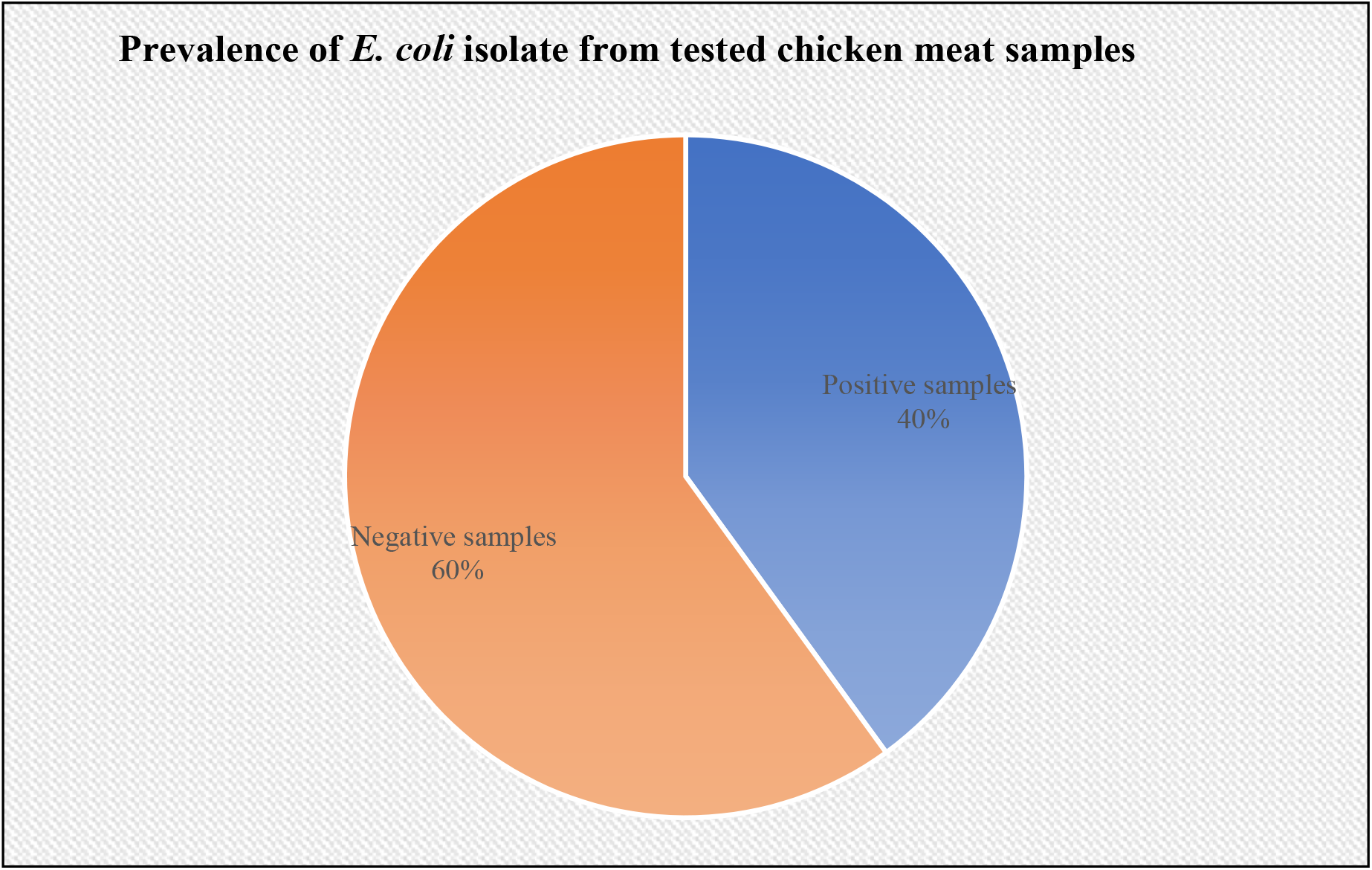
prevalence of E. coli isolate from tested chicken meat samples

**Table 2.** Antimicrobial susceptibility of E. coli strains (n=20) from raw and cooked beef

| Antimicrobial family | Antimicrobial agent | Sensitive |  | Intermediate |  | Resistant |  |
| --- | --- | --- | --- | --- | --- | --- | --- |
|  |  | No | % | No | % | No | % |
| Sulfonamides | Trimethoprim | 20 | 100 | 0 | 00 | 0 | 00 |
|  | Sulfamethoxazole | 16 | 80 | 2 | 10 | 2 | 10 |
| Cephalosporins | Cefepime | 14 | 70 | 4 | 20 | 2 | 10 |
|  | Cefotaxime | 13 | 65 | 4 | 20 | 3 | 15 |
| Tetracycline | Tetracycline | 10 | 50 | 0 | 00 | 10 | 50 |
|  | Doxycycline | 8 | 40 | 2 | 10 | 10 | 50 |
| Quinolones | Ciprofloxacin | 4 | 20 | 4 | 20 | 12 | 60 |
|  | Flumequine | 4 | 20 | 6 | 30 | 10 | 50 |
| Aminoglycosides | Gentamicin | 3 | 15 | 5 | 25 | 12 | 60 |
|  | Streptomycin | 2 | 10 | 5 | 25 | 13 | 65 |
| β-lactame | Penicillin | 2 | 10 | 6 | 30 | 12 | 60 |
|  | Ampicillin | 0 | 00 | 7 | 35 | 13 | 65 |

**Fig 3.**
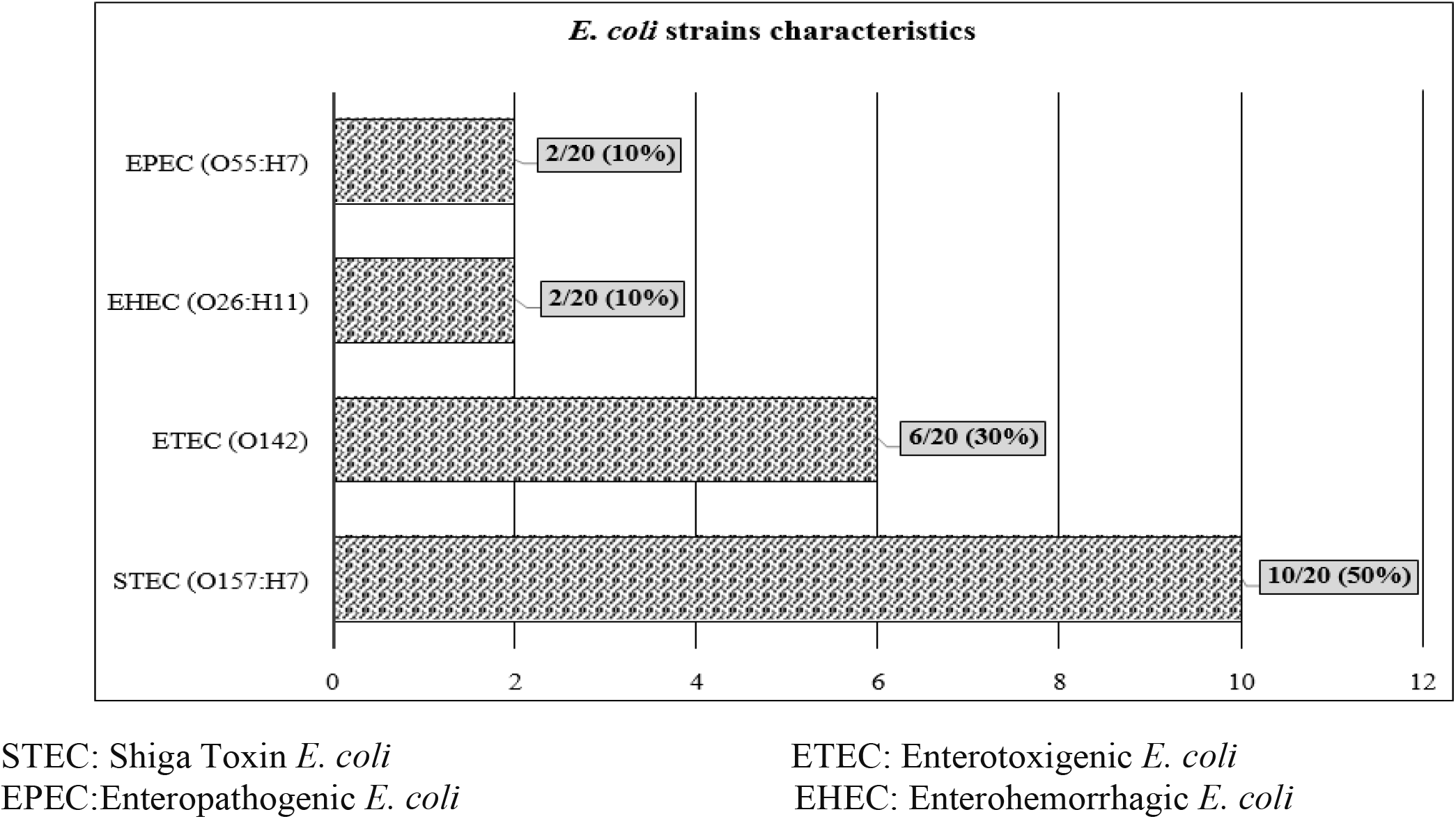
Serotypes of *E. coli* (n=20) isolated from positive chicken samples

**Fig 4.**
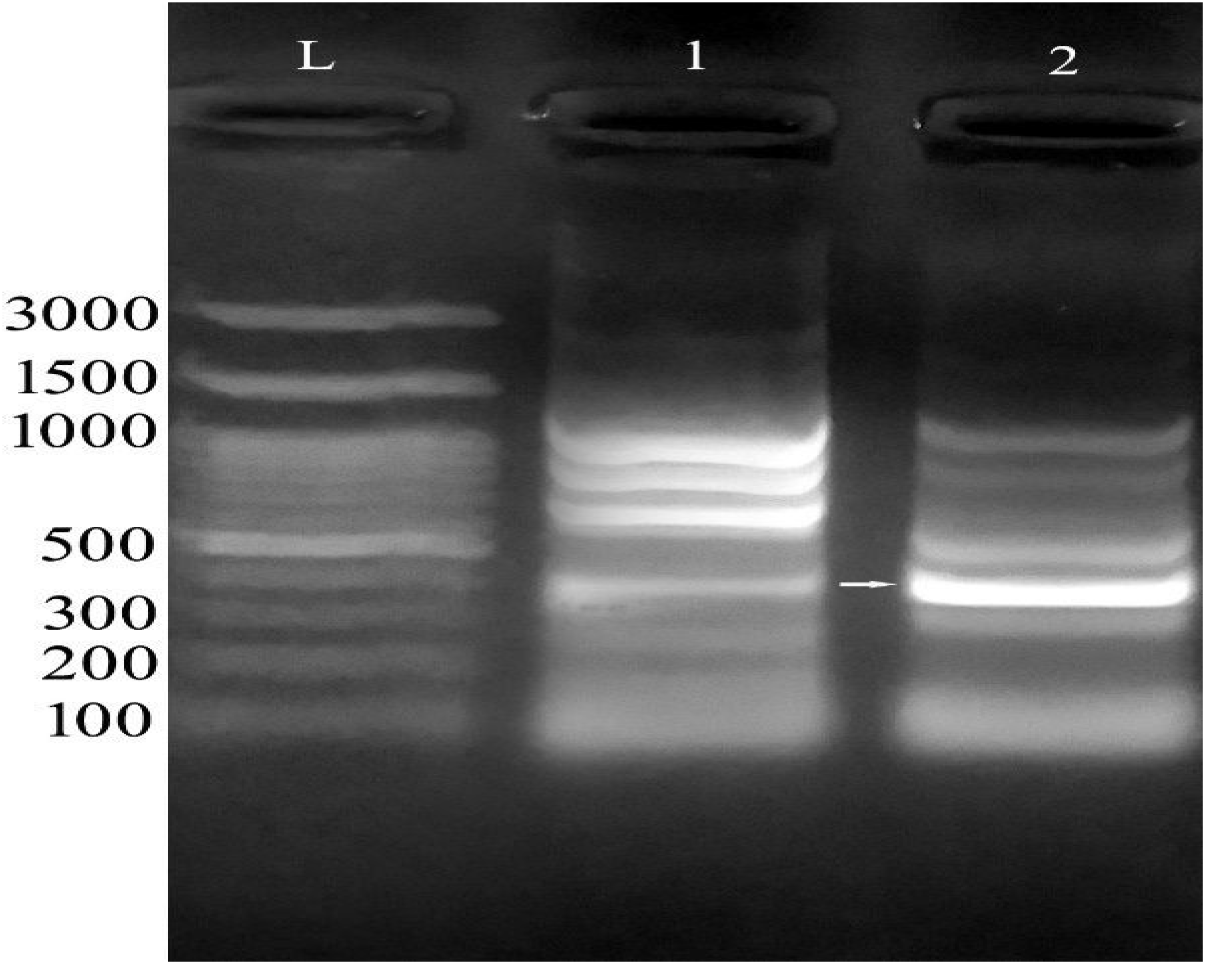
Screening of shiga-like toxin (SLT) – subunit (B) in tested strains. Strain 1 showed a lower density of the amplified gene. Lane (L) represents a standard DNA ladder with 100-3000 bp range

**Fig 5.**
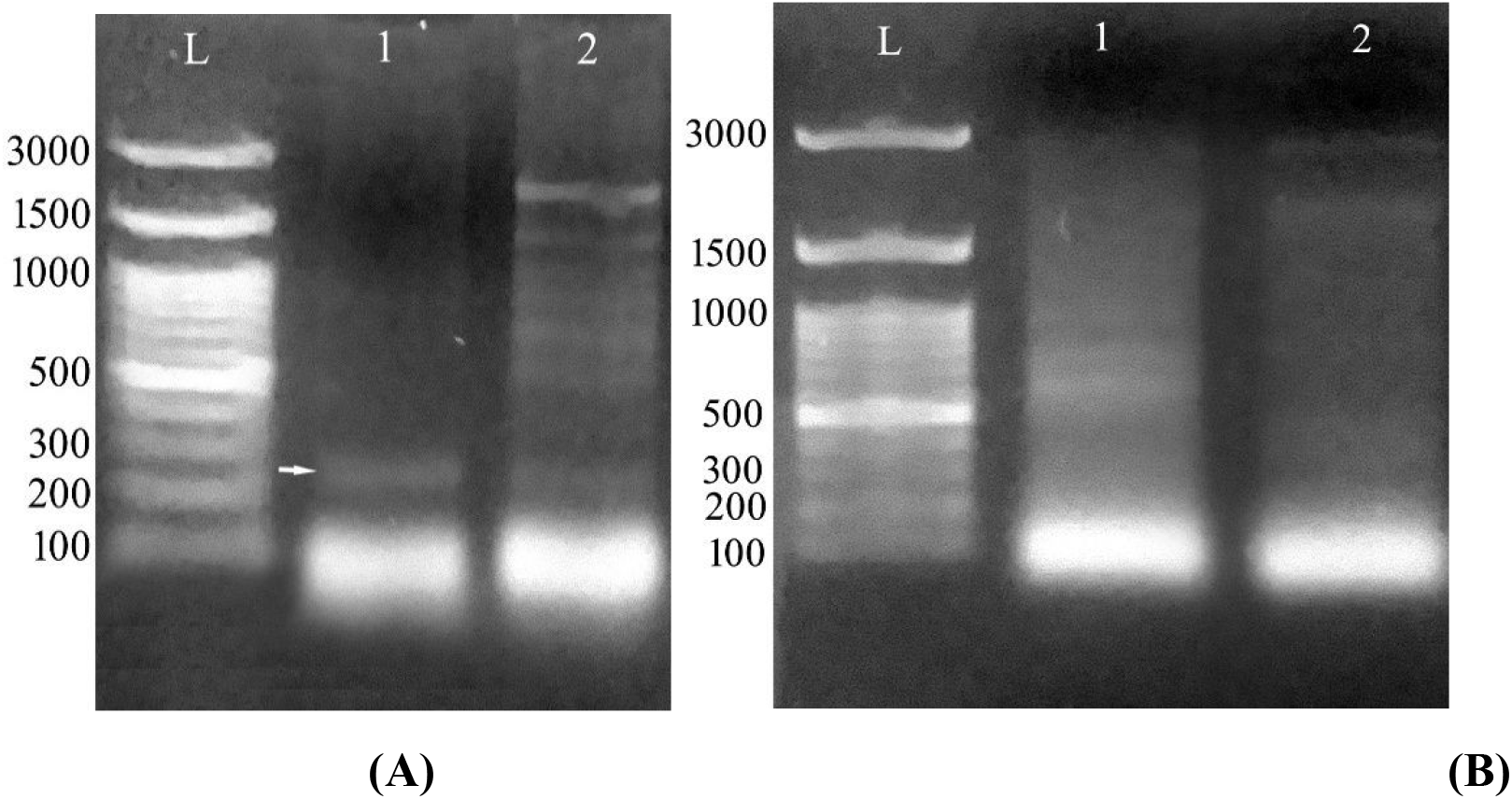
Screening of heat-labile toxin (LT) gene in genomic DNA (A) and plasmid preparations (B). The target gene was detected at a molecular weight of ∼ 200 bp in the genomic DNA of strains (1) and (2).

**Fig 6.**
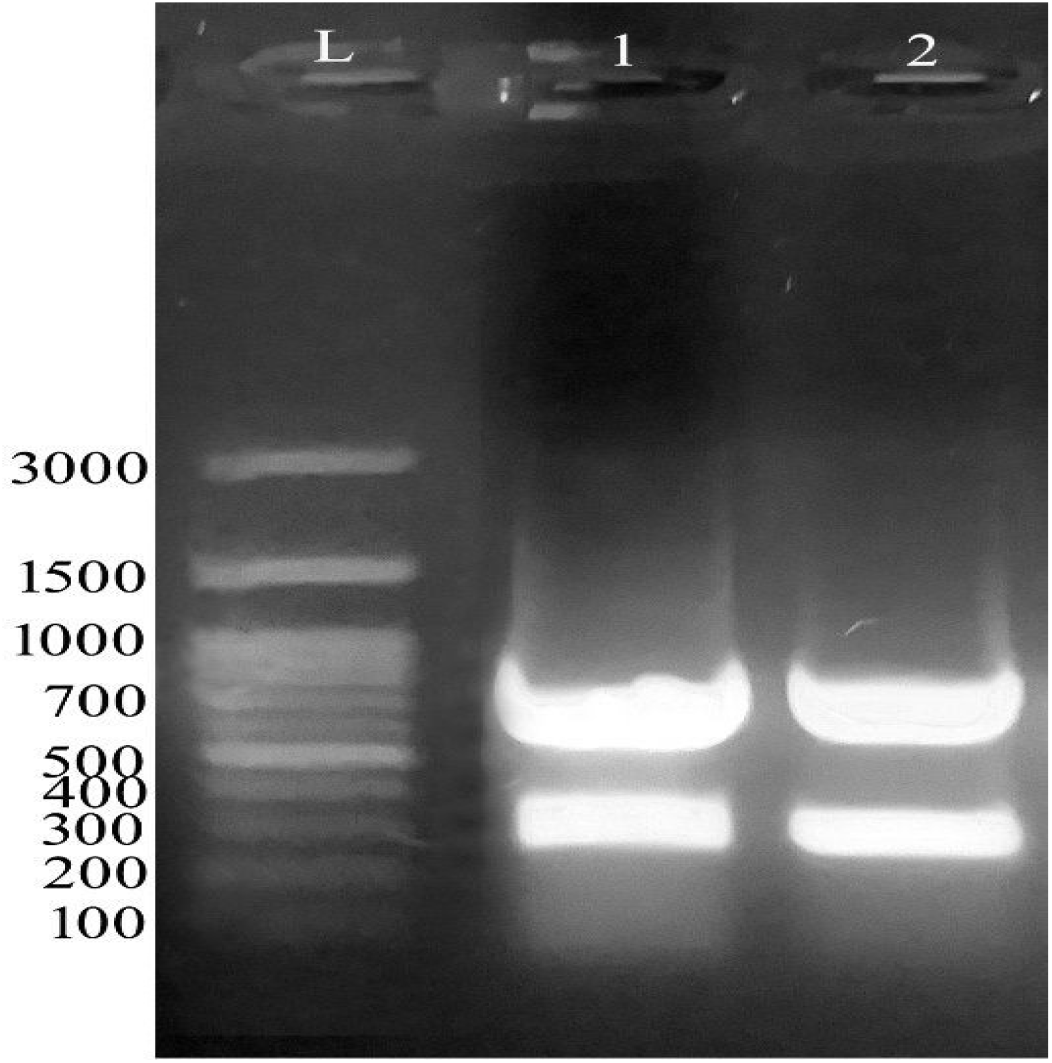
Gentamycin-resistance gene screening: strains screened showed the target fragment of molecular weight ∼ 856 bp corresponding to gentamycin-resistance (aac C2) gene. A minor band with 300 bp was visualized in strains 1 and 2.

**Fig 7.**
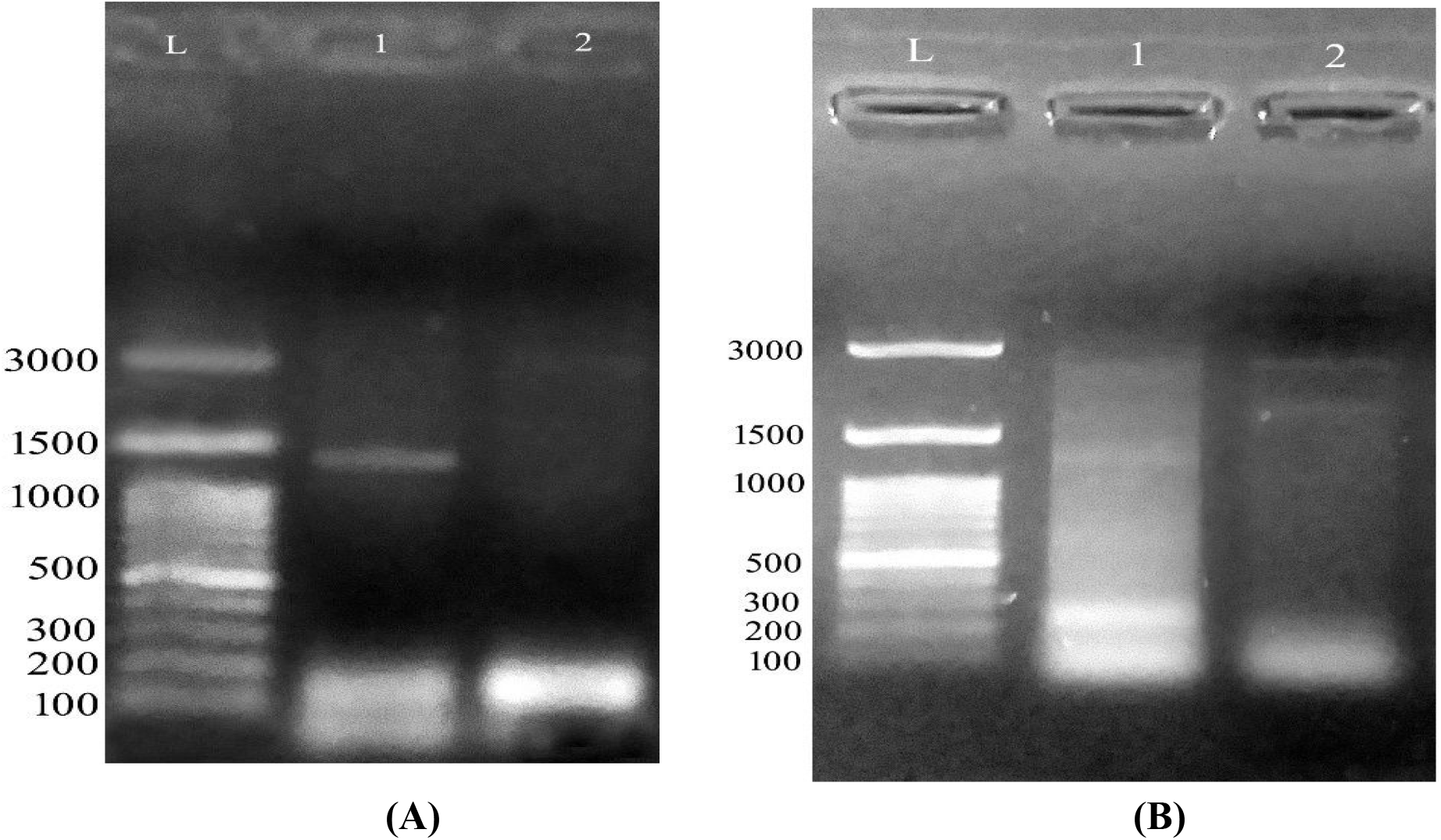
Ciprofloxacin-resistance gene screening: (A) on genomic only strain (1) showed amplification at 1kb size. (B) plasmid.

## Discussion

Concerning bacteriological examination of 100 chicken meat samples, the current study revealed that *E. coli* by 40%. These results agreed with **Partridge, *et al***., **[9]** investigated chicken meat of Mexico and recorded about 35.5%, ***Wu, et. al***., **[10]** who isolated *E. coli* from chicken meat by 35.0%, lower result recorded by **Ngullie, *et al***., **[11]** examined Indian chicken meat and recorded about 31%, **Sato *et al***., **[12]** recorded about 20% in USA chicken meat, **Shaltout, *et. al***., **[13]** who isolated *E*.*coli* from chicken meat hospital meal by 13.33%, **Tomova, *et al***., **[14]** tested Nigerian chicken meat and recorded about 11.1%, **Liu, *et al***., **[15]** found about 10.60% from Croatian chicken meat, **Jakabi, *et. al***., **[16]** who isolated *E. coli* by 9% of from chicken meat. while, **Deng, *et al***., **[17]** detected about 5.92% from the poultry meat in Saudi Arabia, **Schulz, *et al***., **[18]** recorded about 1.56% from a Morocco chicken meat.

This may reflect bad hygienic practice during different stages from slaughtering, handling practices, transportation and excessive handling during preparation of the meal and presence of this microorganism in post processing meat meal indicated that post processing contamination was occurring [19]. *E. coli* is found in the intestinal tract of both humans and animals, finding this organism in ready-to-eat foods is generally viewed as an indication of faecal contamination. Faecal contamination, in turn, indicates that other harmful organisms, whether they be bacterial genera (Salmonella, Shigella, Campylobacter), could be present [20].

Antimicrobial compounds used to avoid and/or treat infections in addition to its benefit as growth promotors in chicken. Its benefit achieved if the antimicrobials selected properly, The antimicrobial susceptibility testing of some *E. coli* strains (n=20) which isolated from chicken meat samples declared that the most effective antimicrobials related to **Sulfonamides**; trimethoprim (20/20) 100%, sulfamethoxazole (16/20) 80%, followed by **Cephalosporins** including; Cefepime (14/20) 70%, Cefotaxime (13/20) 65%, then **Tetracyclines** including; Tetracycline (10/20) 50%, Doxycycline (8/20) 40%, while among tested antimicrobial classes 3 family members recorded weak effect against *E. coli* as following; **Quinolones** including; Ciprofloxacin (4/20) 20%, Flumequine (4/20) 20%, **Aminoglycosides;** Gentamicin 3/20 (15%), Streptomycin (2/20) 10%, and the weakest members related to **β-lactame;** Penicillin 2/20 (10%), Ampicillin (zero%). These results were agreement with **CDC [4]**.

Similar results reported by **Younis *et. al***., **[21] w**ho observed 100% resistance of *E. coli* against penicillin, cefepime 95.8% and amoxicillin 94.5%. according to **Ammar *et. al***., **[22]** many reports declared the resistance of *E. coli* against almost antibiotics due to presence of plasmids genes resistance. **Adeyanju, *et al***., **[23]** and **Bie, *et al***., **[24]** informed that *E. coli* recorded about 90% resistant against tetracycline, ampicillin, trimethoprim-sulphamethozazole, cephalexin, streptomycin, gentamycin. According to **Ramadan *et. al***., **[25]**.

*E. coli* had multi resistance against aminoglycoside, tetracycline, β-lactams, and sulfonamides. **Eid & Erfan [26] and Mohamed, *et al***., **[27]** documented that almost strains of *E. coli* had resistance against β-lactams. According to **Li, *et. al***., **[28]** informed that *E. coli* resisted against; gentamicin, sulfanethazine, sulfadiazine, amoxicillin, tetracycline, ampicillin, chloramphenicol, and ceftriaxone. **Zhang, *et. al***., **[29]** found that about 60% of E. coli which isolated from food of animal origin were resistant against fluoroquinolone. **Tang, *et. al***., **[30]** noticed that about 36.8%, 35.0% and 34.1% resisted norfloxacin, ciprofloxacin and enrofloxacin.

Twenty *E. coli* isolates were serologically 10/20 (50%) isolates were typed as STEC (O157:H7), 6/20 (30%) isolates were ETEC (O142), 2/20 (10%) isolate was EHEC (O26:H11). Finally, 2/20 (10%) of isolates were typed as EPEC (O55:H7). **Shiga-Like Toxin (SLT) Screening documented as following:** the subunit B of shiga-like toxin (SLT) gene appeared as a homogenous band in their genomic DNA at 300 bp molecular weight.

Results declared that strain (1) were the lowest amplification detected comparing to strain **Heat-Labile Toxin (LT) Screening declared that:** Heat-labile toxin (LT) gene was screened in both genomic DNA and plasmid preps in tested strains. A fragment of ∼ 200 bp was detected in both strains (1) and (2). **Gentamycin Resistance Screening observed:** Gentamycin resistance gene (aac C2) fragment was detected in strains as a fragment of molecular weight ∼ 856 bp but with a minor band of molecular weight of nearly 300 bp. **Ciprofloxacin Resistance Screening viewed that:** Ciprofloxacin-resistance gene was also screened in both genomic and plasmid preparations in strains under test. A sharp band at 1 kb was clearly detected in strain (1) while it was absent in strain (2). In plasmid preps, no amplification of the target gene could be detected in strains tested. These results agreed with the obtained results by **Momtaz and Jamshidi, [31]** who isolated O serogroups, especially O1, O35, O2, O15, O8, O18, O88, O78, O109, and O115. Ćwiek, et al., (2021) informed that there were two serotypes which is very similar to eae genes of E. coli serotypes O55:H7 and O157:H7 strains, eaeA gene and enteropathogenic E. coli and EHEC. Furthermore, Li, et. al., (2020) reported that E. coli lose SLT genes which leading to false-negative results. In addition to the screening of EHEC eaeA gene declared the positive results of *E. coli* not O157:H7. While HEC O157 detected by 60-MDa plasmid. Kim, et. al., (2019) observed that gene (SLT I, II and eaeA), which indicated the presence of EHEC O157 strains. Kluytmans-van, et. al., (2016) documented that Stx gene used for detection of EHEC strains they added that the virulence genes contained extraintestinal infections genes such as; (cdt2, afaD8, cdt3, bmaE, iroN, iucD, iutA and traT). **Villegas, *et.al***., **[32] reported that** etpD gene incate the presence of ETEC strains. While they detected enterohemorrhagic *E. coli* strains O157:H7 by RIMD 0509952 and EDL933. Presence of fimH gene indicated presence of non-pathogenic *E. coli*, ETEC, EPEC, and UPEC strains.

## Conclusion

In summary, the research detected the virulence genes of E. coli from chicken meat including different somatic capsular & antigen genes. It is very important to control STEC as it represents a danger to the poultry consumers. *E. coli* is naturally found in our daily diet and hazarded the food biosafety and public health. We recommended to increase the hygienic measures during slaughtering, processing and/or handling of chicken carcasses. The study recommended also by avoidance the unnecessary usage of any antimicrobial compounds to life chicken and human to avoid appearance of new antimicrobials resistant.

## Acknowledgement

This work was funded by the University of Jeddah, Jeddah, Saudi Arabia, under grant No. (UJ-23385-DR). The authors, therefore, acknowledge with thanks the University of Jeddah technical and financial support.

